# MCT4 is a high affinity transporter capable of exporting lactate in high-lactate environments

**DOI:** 10.1101/586966

**Authors:** Y Contreras-Baeza, PY Sandoval, R Alarcón, A Galaz, F Cortés-Molina, K Alegría, F Baeza-Lehnert, R Arce-Molina, A Guequén, CA Flores, A San Martín, LF Barros

**Affiliations:** Centro de Estudios Científicos - CECs, Valdivia, Chile.; Universidad Austral de Chile, Valdivia, Chile.

## Abstract

MCT4 is an H^+^-coupled transporter expressed in metastatic cancer cells, macrophages, and other highly glycolytic cells, where it extrudes excess lactate generated by the Warburg phenomenon or by hypoxia. Intriguingly, its reported K_m_ for lactate, obtained with pH-sensitive probes, is more than an order of magnitude higher than physiological lactate. Here we examined MCT4-rich MDA-MB-231 cells using the FRET sensor Laconic and found a median K_m_ for lactate uptake of only 1.7 mM, while parallel estimation in the same cells with a pH probe gave a K_m_ of 27 mM. The median K_m_ of MCT4 for lactate was 0.7 mM in MCT4-expressing HEK293 cells and 1.2 mM in human macrophages, suggesting that high substrate affinity is a robust property of the transporter. Probed with the FRET sensor Pyronic, MCT4 showed a K_m_ for pyruvate of only 4.2 mM in MDA-MB-231 cells, as opposed to > 150 mM reported previously. We conclude that prior estimates of MCT4 affinity based on pH probes were severely biased by the confounding action of pH regulatory mechanisms. Numerical simulation showed that MCT4, but not MCT1 or MCT2, endows cells with the capability of lactate extrusion in high lactate environments. The revised kinetic properties and novel transport assays may help in developing small-molecule MCT4 blockers for research and therapy.

## INTRODUCTION

Cancer cells ferment glucose to lactate in the presence of oxygen, a phenomenon originally described by Otto Warburg and colleagues in the 1920s and later found to promote tumor growth and malignancy ^1–4^. In addition to fostering glycolysis by end-product removal, cytosolic alkalinization and NAD^+^ recycling, the co-extrusion of lactate and protons causes interstitial acidification, which along lactate itself favors tumor invasiveness and facilitates immune response evasion ^5^. Lactate levels were double in cervical tumors with metastatic spread compared with malignancies in patients without metastases ^6^. Lactic acid release is also a physiological process, as in exercising skeletal muscle, in active macrophages and in brain astrocytes ^7–9^. The transport of lactate in most mammalian cells is mediated by members of the SLC16A family of H^+^-coupled monocarboxylate transporters (MCTs) ^10^. Malignant tumors overexpress MCT1 (SLC16A1) and/or MCT4 (SLC16A3), the latter being characteristic of metastatic cancer in association with HIF-1α upregulation ^11^. Potent small-molecule inhibitors specific for MCT1-2 have been synthetized, one of which is currently undergoing a Phase I clinical trial ^12^. However, the development of MCT4-specific drugs is lagging.

MCT4-endowed cells, both healthy and cancerous, are the strongest lactate producers. So it seems puzzling that, at about 30 mM ^10, 13, 14^, the K_m_ of MCT4 for lactate is more than one order of magnitude higher than the levels of lactate prevailing in tissues and even within hypoxic tumors ^6,15^. Taken at face value, this means that MCT4 runs at a small fraction of its capacity. In contrast, MCT1 has a K_m_ of 3-5 mM ^10^. Kinetic transport parameters are determined by measuring initial rates of uptake at increasing concentrations of radiolabeled substrate. Unfortunately, this is not practical for MCTs in mammalian cells, because uptake is too fast, demanding high levels of radioactivity and sophisticated stop-flow devices. The introduction of pH-sensitive dyes in the 1980s revolutionized the field by permitting MCT activity determinations with high spatiotemporal resolution ^16, 17^. With the additional advantage that any substrate could be studied with the same probe, most of what we know about the functionality of the monocarboxylate transporters was learnt from substrate-induced acidification. However, there was a caveat. In order to obtain detectable acidifications, experiments had to be done in the absence of bicarbonate.

Genetically-encoded FRET nanosensors have been recently used by several laboratories to directly monitor lactate and pyruvate dynamics in various cell types, *in vitro* and *in vivo* ^18–27^. During the characterization of a MCT4-rich cell line with a FRET sensor we detected robust transport at low lactate concentrations. The present manuscript describes a set of experiments prompted by that observation.

## METHODS

Standard reagents and inhibitors were acquired from Sigma or Merck. Plasmids encoding the sensors Laconic ^18^ and Pyronic ^19^ are available from Addgene (www.addgene.org). Ad Laconic and Ad Pyronic (serotype 5) were custom made by Vector Biolabs.

### Cell culture

MDA-MB-231 cells were acquired from the American Type Culture Collection (ATCC) and cultured at 37 °C without CO_2_ in Leibovitz medium (ThermoFisher). Cultures were transfected at 60% confluence using Lipofectamine 3000 (ThermoFisher) or alternatively, exposed to 5 x 10^6^ PFU of Ad Laconic or Ad Pyronic and studied after 24-72 h. The generation of the MDA-LAC cell line will be described elsewhere (Mito Toxy reporter; Contreras et al, submitted doi: https://doi.org/10.1101/583096). HEK293 cells were acquired from the ATCC and cultured at 37 °C in 95% air/5% CO_2_ in DMEM/F12 supplemented with 10% fetal bovine serum. Cultures were transfected at 60% confluence using Lipofectamine 3000 (ThermoFisher) and studied after 24-72 To obtain macrophages, blood was collected by venepuncture from ten healthy male volunteers. Age of donors ranged from 25 to 45 years. Ethical guidelines stipulated by the Declaration of Helsinki principles were adhered to. Approval was obtained from the Medical Ethical Committee of the Faculty of Medicine, Universidad Austral de Chile. All donors were informed about the nature of the studies and gave their written consent to participate. Samples were treated anonymously. Monocytes were isolated from whole blood treated with 3.8% sodium citrate, by Percoll™ density gradient centrifugation (GE Healthcare, Sweden). Macrophage differentiation of human monocytes was induced by treatment with 25 nM macrophage colony-stimulating factor for seven days. Human monocytes-derived macrophages were treated with 100 ng/mL IFN-γ and 10 ng/mL lipopolysaccharide (LPS) for M1 differentiation for 48 h. After isolation cells were maintained in RPMI media supplemented with 10% fetal bovine serum and 1% pyruvate. Cytokines were obtained from PeproTech (USA), LPS (from *Pseudomonas aeruginosa*) was from Sigma-Aldrich (USA). For lactate measurements, macrophages maintained in culture for 6 to 7 days were incubated for 24 h with 7×10^10^ PFU Ad Laconic and imaged 6-7 days later.

### Imaging

Cells were imaged at 35 °C in a 95% air/5% CO_2_-gassed KRH-bicarbonate buffer of the following composition (in mM): 112 NaCl, 3mM KCl, 1.25 CaCl_2_, 1.25 MgSO_4_, 10 HEPES, 24 NaHCO_3_, pH 7.4. Alternatively, NaHCO_3_ was equimolarly replaced with NaCl. Glucose, lactate, pyruvate and inhibitors were added as indicated in the figure legends. Laconic and Pyronic were imaged using an upright Olympus FV1000 confocal microscope equipped a 20X water immersion objective (NA 1.0). Laconic and Pyronic were imaged at 440 nm excitation/480 ± 15 nm (mTFP) and 550 ± 15 (Venus) emissions. BCECF was ester-loaded at 0.1 μM for 3-4 min and the signal was calibrated by exposing the cultures to solutions of different pH after permeabilizing the cells with 10 μg/ml nigericin and 20 μg/ml gramicidin in an intracellular buffer. BCECF was sequentially excited at 440 and 490 nm (0.05 s) and imaged at 535/30 nm using an Olympus BX51 microscope (20X water immersion objective, NA 0.95) equipped with a CAIRN monochromator and Optosplit II (Faversham, UK) and a Hamamatsu Rollera camera.

### Immunodetection

For immunoblotting cultured cells were scraped into cold phosphate-buffered saline (PBS 1X) followed by centrifugation at 5,000 rpm for 5 min at 4°C. The cell pellet was then suspended in cold RIPA 1X (radioimmune precipitation assay) lysis buffer (50 mM Tris-HCl, pH 7.4, 150 mM NaCl, 0.1% SDS, 0.5% sodium deoxycholate, 1% Nonidet P40, 10 mM N-ethylmaleimide, 0.1 mM phenylmethylsulphonyl fluoride, 1 µg/ml aprotinin, 1 µg/ml leupeptin, and 1 µg/ml pestatin A). After 30 min on ice, unlysed cells and nuclei were pelletted at 12,000 rpm for 15 min at 4°C. The protein concentration of the supernatant was determined by Bio-Rad Dc Protein Assay using BSA standards. Protein samples (50 µg) were loaded onto 10% (w/v) SDS-polyacrylamide gels and electrotransferred onto nitrocelulose membranes. Antibodies used were rabbit anti-MCT4 (1:250; AB3314P, Merck Millipore), mouse anti- β-actin (1:2,000; 8H10D10, Cell Signaling), peroxidase-conjugated antirabbit (1:20,000; 111-035-144, Jackson) and peroxidase-conjugated antimouse (1:20,000; #7076, Cell Signaling). Signals were revealed using a chemiluminescence kit (SuperSignal(tm) West Femto, ThermoFisher), following the instructions of the manufacturer. For immunocytochemistry macrophages fixed with 4% paraformaldehyde and permeabilized with 0.1% PBS-Triton X-100 were blocked with 3% Bovine Serum Albumin (BSA) plus 10% Normal Goat Serum (Vector Laboratories). Samples were probed with a primary antibody against MCT4 (1:1000; AB3314P, Merck Millipore and AlexaFluor488 secondary antibody (1:2000; A11008, Invitrogen). Nuclei were stained with 4,6-diamidino-2-phenylindole (1:50,000; D3571, Invitrogen). Samples were embedded into fluorescence mounting medium (Dako) and analyzed by confocal microscopy (Leica DMi8, Leica Microsystems) using a 63x oil objective. Image analysis was performed using Leica and ImageJ software.

### Mathematical modeling

Cellular lactate and pyruvate dynamics were simulated using Berkeley Madonna software and the following set of ordinary differential equations,

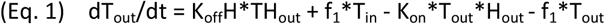

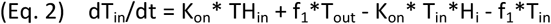

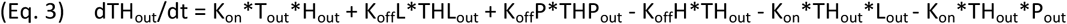

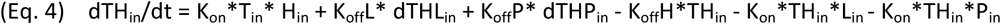

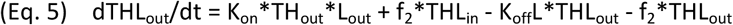

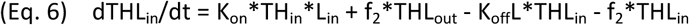

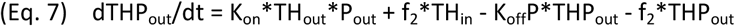

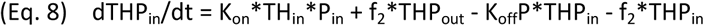

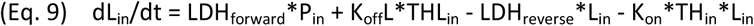

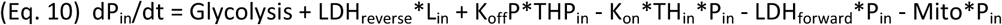

where equations 1 to 8 represent the eight possible conformations of the MCT carrier: outward-and inward-facing, either empty (T_out_ and T_in_), loaded with a proton (TH_out_ and TH_in_), loaded with both proton and lactate (THL_out_ and THL_in_) and loaded with both proton and pyruvate (THP_out_ and THP_in_). Equations 9 and 10 represent cytosolic lactate and pyruvate. The association constant K_on_ for protons, lactate and pyruvate was set at 10^8^ M^-1^ *s^-1^ for the three isoforms (diffusion-limited); respective dissociation constants K_off_H, K_off_L and K_off_P were 20 s^-1^, 7.6*10^7^ s^-1^ and 7.6*10^6^ s^-1^ for MCT1, 20 s^-1^, 7.6*10^6^ s^-1^ and 7.6*10^5^ s^-1^ for MCT2, and 20 s^-1^, 2.4*10^7^ s^-1^ and 6.4*10^7^ s^-1^ for MCT4. Carrier translocation rates f_1_ (empty) and f_2_ (loaded) were set at 200 s^-1^ and 3000 s^-1^. Rate constants were 0.01 s^-1^ (Mito, mitochondrial pyruvate import), 0.5 s^-1^ (LDH_forward_, pyruvate to lactate) and 0.025 s^-1^ (LDH_reverse_, lactate to pyruvate). With these parameters and cytosolic and extracellular pH values of 7.2 (63 nM) and 7.4 (40 nM), apparent zero-trans K_m_ (K_zt_) values for lactate and pyruvate uptake were: 5 and 0.5 mM for MCT1, 0.5 and 0.05 mM for MCT2, and 1.7 and 4.2 mM for MCT4.

### Statistical Analysis

Statistical analyses were carried out with SigmaPlot software (Jandel). For normally distributed variables, differences were assessed with the Student’s t-test (pairs) and with ANOVA followed by the Tukey-Kramer *ad hoc* test (groups). For variables that failed the normality test, differences were assessed with the Kruskal-Wallis one way ANOVA on ranks (groups). *, p < 0.05; ns (non-significant), p > 0.05. The number of experiments and cells is detailed in each figure legend.

## RESULTS

### MCT4 mediates monocarboxylate transport in MDA-MB-231 cells

To study the functional properties of MCT4, we expressed the genetically-encoded FRET lactate sensor Laconic ^18^ in MDA-MB-231 cells, a human breast cancer cell line conspicuous for its high levels of MCT4 ^28, 29^. Fig. 1A shows MDA-LAC, a cell line generated with MDA-MB-231 cells stably expressing Laconic. Fig. 1B shows that the abundance of MCT4 in these cells is almost as high as that achieved by overexpressing MCT4 in HEK293 cells under the strong CMV promoter, and that MCT4 levels are not diminished by expression of the FRET sensor. Exposure of MDA-MB-231 cells to a lactate load caused a rapid increase in intracellular lactate, demonstrative of high permeability (Fig. 1C). The functionality of MCT4 was tested by pharmacological means. As there are no commercially available inhibitors specific for MCT4, we tested compounds of overlapping selectivity. pCMBS, which inhibits MCT1 and MCT4 but not MCT2 ^30^, caused lactate accumulation (Fig. 1C). In contrast AR-C155858 ^31^, which blocks MCT1 and MCT2 but not MCT4, had no effect (Fig. 1D). Taken together, these results show that the tonic export of lactate by MDA-MB-231 cells is mediated by MCT4. The insensitivity to AR-C155858 is in agreement with the reported absence of detectable MCT1 expression in these cells ^28, 29^. Next, the effect of the pharmacological inhibitors was tested on lactate uptake. Consistent with the efflux data, pCMBS blocked the uptake of lactate while AR-C155858 did not (Figs. 2A-B). Moreover, diclofenac, a structurally-unrelated MCT1 & MCT4 blocker ^32^, also abrogated the influx of lactate whereas AZD3965 ^33^, a structurally-unrelated blocker of MCT1 & MCT2 (but not of MCT4), had no effect (Figs. 2C-D). These results provide functional confirmation that MCT4 is responsible for the bidirectional transport of lactate across the plasma membrane of MDA-MB-231 cells.

**Figure 1.**
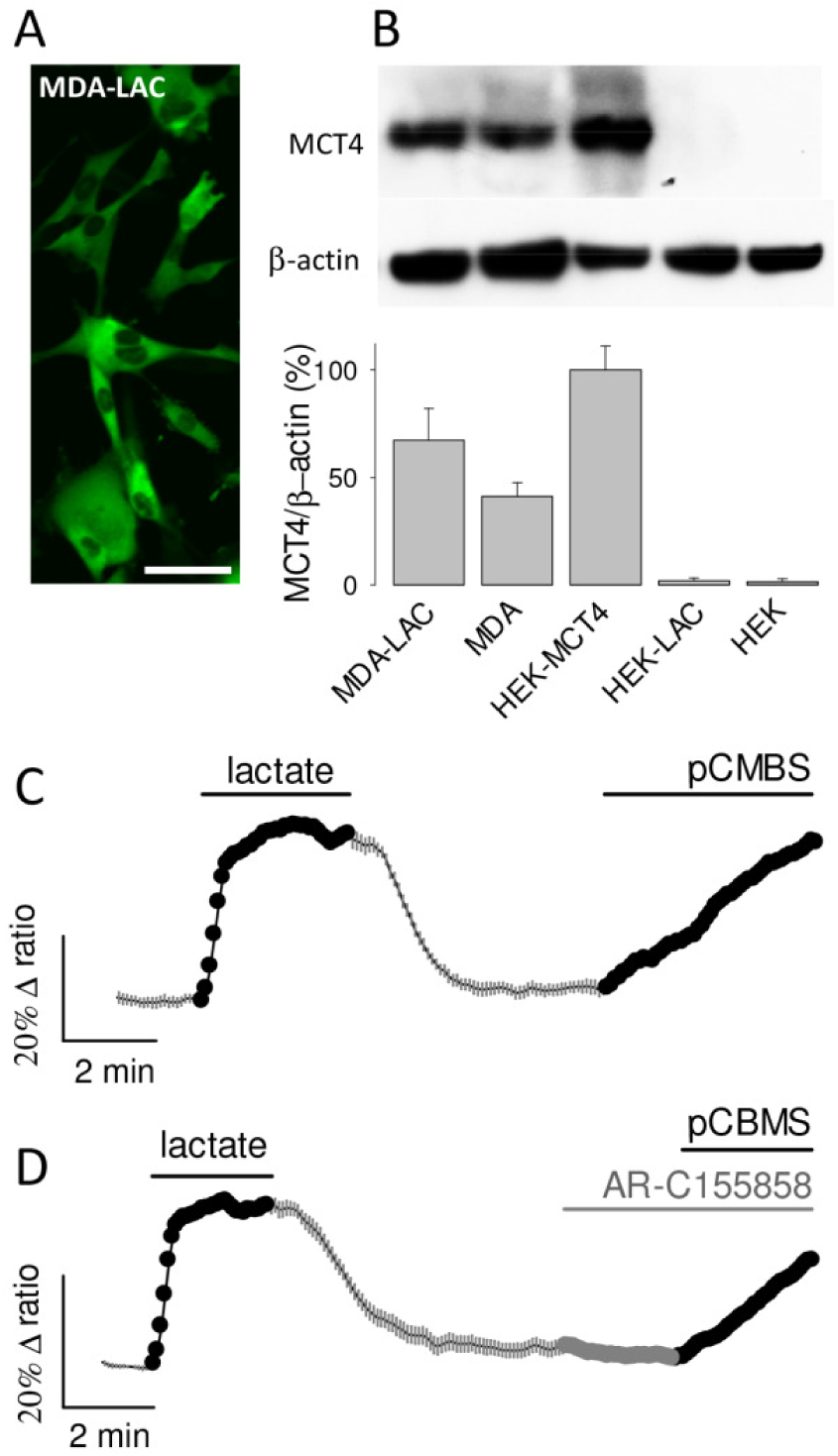
MCT4 mediates tonic lactate efflux in MDA-MB-231 cells. (A) MDA-MB-231 cells permanently expressing the lactate sensor Laconic (MDA-LAC). Bar represents 50 μm. (B) Immunodetection of MCT4 in extracts from: MDA-LAC, wild type MDA-MB-231 cells (MDA), HEK293 cells transiently expressing MCT4 (HEK-MCT4), HEK293 cells transiently expressing Laconic (HEK-LAC) and wild type HEK293 cells (HEK). Bar graphs show mean ± SEM (3 separate preparations). (C-D) MDA-MB-231 cells expressing Laconic were exposed to 10 mM lactate and then to 1 μM AR-C155858 and/or 250 μM pCMBS as indicated, mean ± SEM (10 cells from single experiments, representative of three experiments for each protocol).

**Figure 2.**
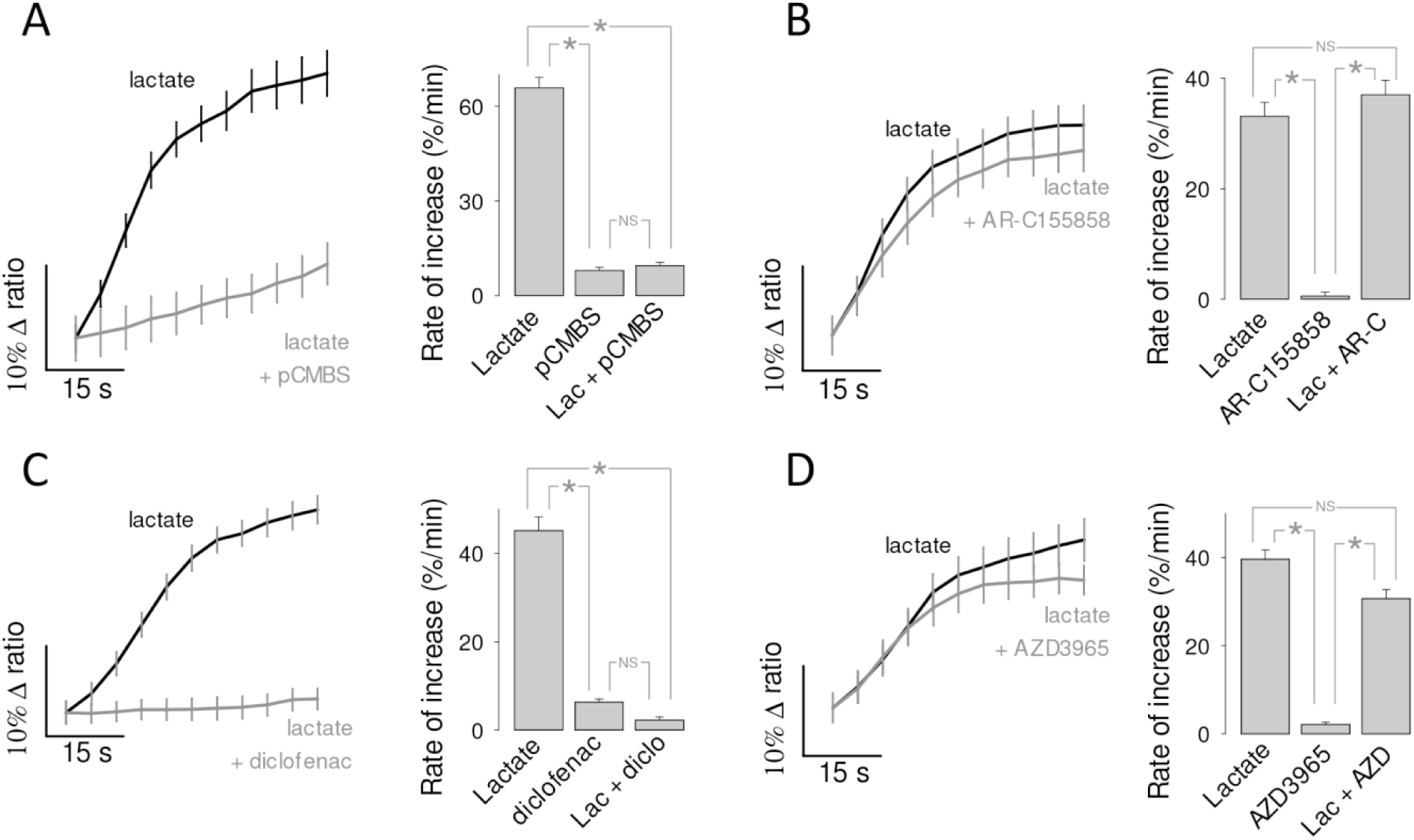
MCT4 mediates the influx of lactate in MDA-MB-231 cells. The uptake of 10 mM lactate by MDA-LAC cells was monitored before and during exposure to MCT inhibitors. Bar graphs show the initial rates of uptake and the rate of accumulation elicited by the inhibitor itself. (A) 250 μM pCBMS, inhibits MCT1 and MCT4. (B) 1 μM AR-C155858, inhibits MCT1 and MCT2. (C) 1mM diclofenac, inhibits MCT1 and MCT4. (D) 10 μM AZD3965, inhibits MCT1 and MCT2.

### MCT4 of MDA-MB-231 cells is a high affinity lactate/pyruvate transporter

The affinity of MCT4 for lactate was determined by exposing MDA-MB-231 cells expressing the FRET lactate sensor to an increasing extracellular concentration of lactate, as described previously in MCT1-expressing cardiomyocytes ^20^. Control uptakes were included at the beginning and end of the protocol to ensure that measurements were reproducible (data not shown). As illustrated in Fig. 3A, robust lactate uptake was already apparent at low lactate concentrations. Plotting uptake rates against lactate concentration revealed K_m_ values in the low milimolar range (Fig. 3B). To investigate possible confounding effects of experimental conditions, the protocol was repeated in the presence and absence of bicarbonate, at 23 °C and at 35 °C, and in the absence and presence of AR-C155858 (to eliminate possible minor contributions of MCT1 and MCT2), in cells in which Laconic was expressed by transfection, an adenoviral vector, or in a stable cell line. As no strong differences under these experimental conditions were detected, the data were pooled together. The median K_m_ for the uptake of lactate was 1.7 mM. In view of this inordinate high affinity for lactate, an analogous experimental approach was applied to characterize the transport of pyruvate, using the FRET sensor Pyronic ^19^. Reportedly, the affinity of MCT4 for pyruvate is so low that it lies beyond the measurable range (> 150 mM ^34^). However, we obtained a median K_m_ of 4.2 mM (Fig. 4). Thus, the affinity of MCT4 for lactate and pyruvate in MDA-MB-231 cells was found to be over one order of magnitude higher than anticipated.

**Figure 3.**
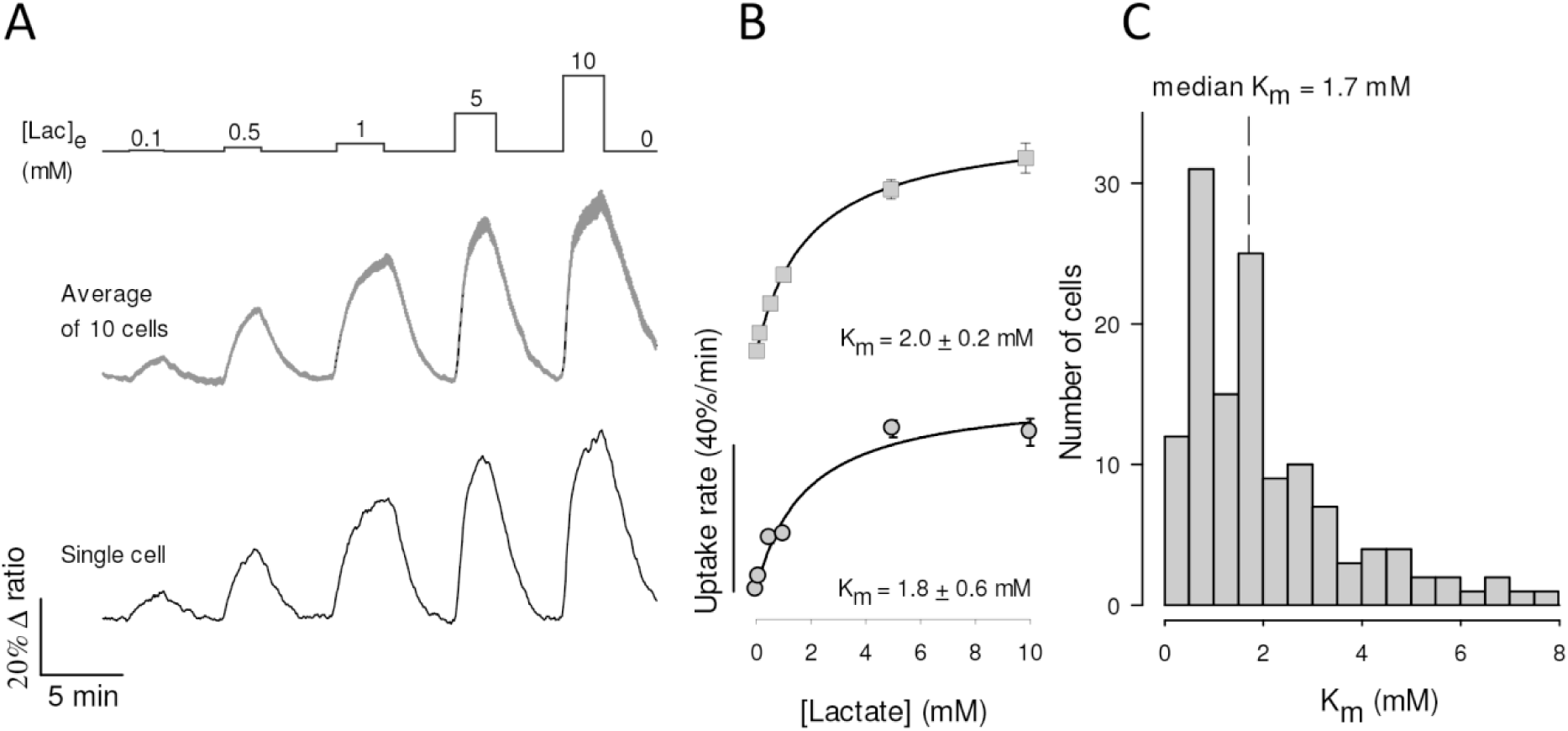
High affinity lactate transport in MDA-MB-231 cells. (A) MDA-LAC cells were exposed to increasing concentrations of lactate, from 0.1 to 10 mM, as indicated. Responses of intracellular lactate in a single cell (bottom) and in 10 cells from a representative experiment (mean ± SEM, top) are shown. (B) Dose-response of the initial rate of lactate uptake, from the same cells depicted in A. Mean ± SEM. K_m_ values were obtained by fitting a rectangular hyperbola to the data. (C) Frequency distribution of K_m_ determinations from 133 cells in ten experiments. Median K_m_ was 1.7 mM.

**Figure 4.**
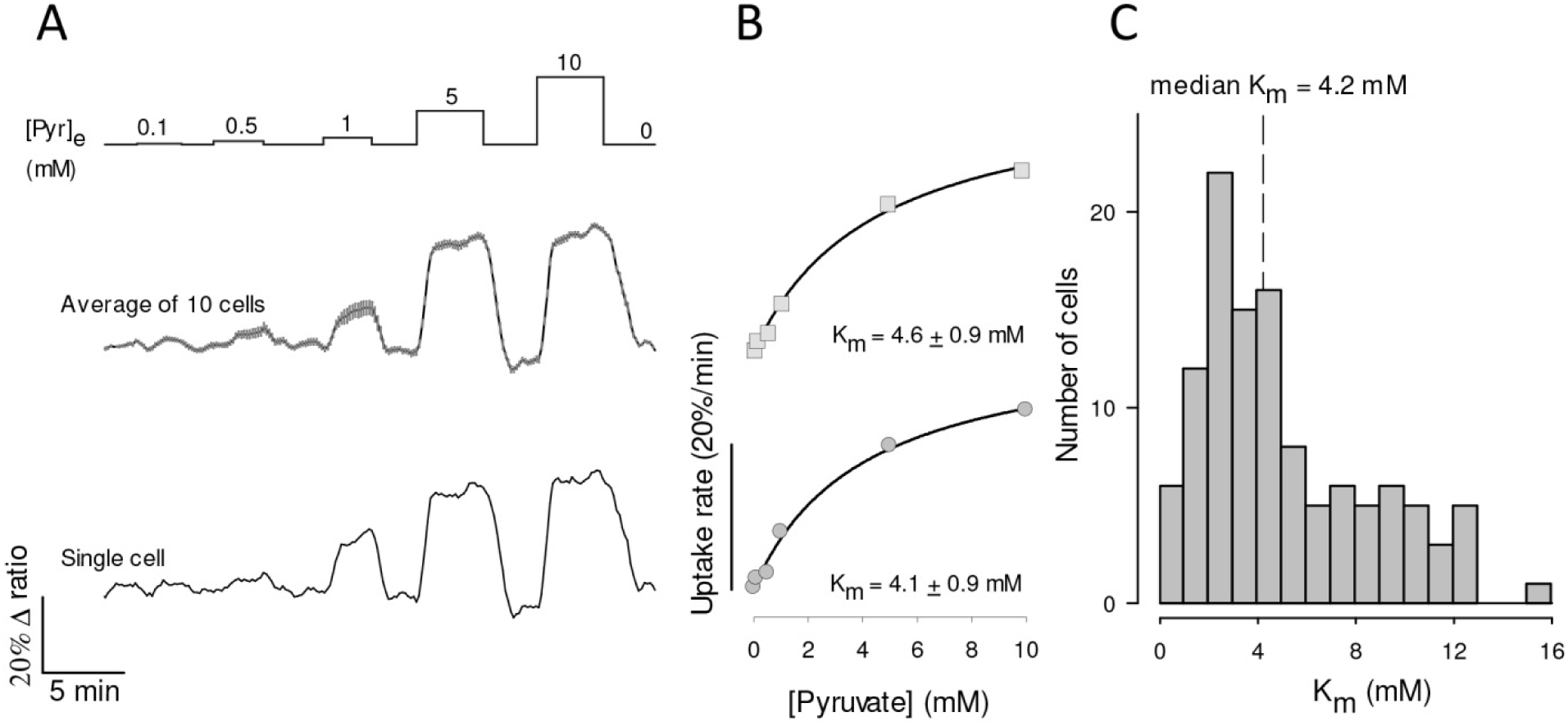
High affinity pyruvate uptake in MDA-MB-231 cells. (A) MDA-MB-231 cells expressing pyronic were exposed to increasing concentrations of pyruvate, from 0.1 to 10 mM, as indicated. The responses of a single cell (bottom) and of 10 cells from a single experiment (mean ± SEM, top) are shown. (B) Dose-response of the initial rate of pyruvate uptake, from the same cells depicted in A. Mean ± SEM. K_m_ values were obtained by fitting a rectangular hyperbola to the data. (C) Frequency distribution of K_m_ determinations from 117 cells in thirteen experiments. The median K_m_ in this series was 4.2 mM.

### Bicarbonate interferes with pH estimation of MCT4 activity

To evaluate MCT4 activity from its effects on intracellular pH, MDA-MB-231 cells were loaded with the pH-sensitive dye BCECF. Exposure to lactate did acidify the cells, but in contrast with the accumulation of lactate measured directly with the FRET sensor, the acidification was highly sensitive to bicarbonate (Fig. 5A). In bicarbonate, the rate of acidification induced by lactate did not show saturation. On the contrary, it jumped by a factor of 8 between lactate exposure of 10 and 20 mM. This non-linear behavior suggests that at 20 mM lactate the flux via MCT4 surpassed the capacity of the cells to muffle protons (Fig. 5B). When bicarbonate was replaced with the impermeant buffer HEPES (no HCO_3_), intracellular pH became more sensitive to lactate challenges and some degree of saturation appeared (Fig. 5A-B). A median K_m_ of 27 mM (26 cells three experiments) could be estimated, which is not deemed accurate as it lies beyond the highest lactate concentration applied. Still, this high K_m_ is in agreement with previous determinations of MCT4 lactate affinity in several cell types using pH, which range from 30 to 40 mM ^34, 35^. When BCECF-loaded MDA-MB-231 cell were challenged with pyruvate the results were similar: insensitivity in the presence of bicarbonate and responses being detected in the absence of bicarbonate only at > 5 mM pyruvate (Fig. 5B). There was no apparent saturation of the rate of acidification with or without bicarbonate, so that K_m_ values could not be estimated, as reported ^34,35^. Of note, bicarbonate omission should not be expected to eliminate the problem of buffering, because bicarbonate represents a minor fraction of the buffering power of mammalian cells ^36, 37^. In addition to buffering, mammalian cells possess efficient systems for the extrusion of protons, including carbonic anhydrase, Na^+^/H^+^ exchangers and Na^+^/bicarbonate cotransporters, some of which have recently been found strategically located in the vicinity of MCTs ^38^. We conclude that buffering/muffling reduces the impact of MCT4-mediated proton transport on intracellular pH, particularly at low substrate concentrations, introducing a bias in the determination of affinity.

**Figure 5.**
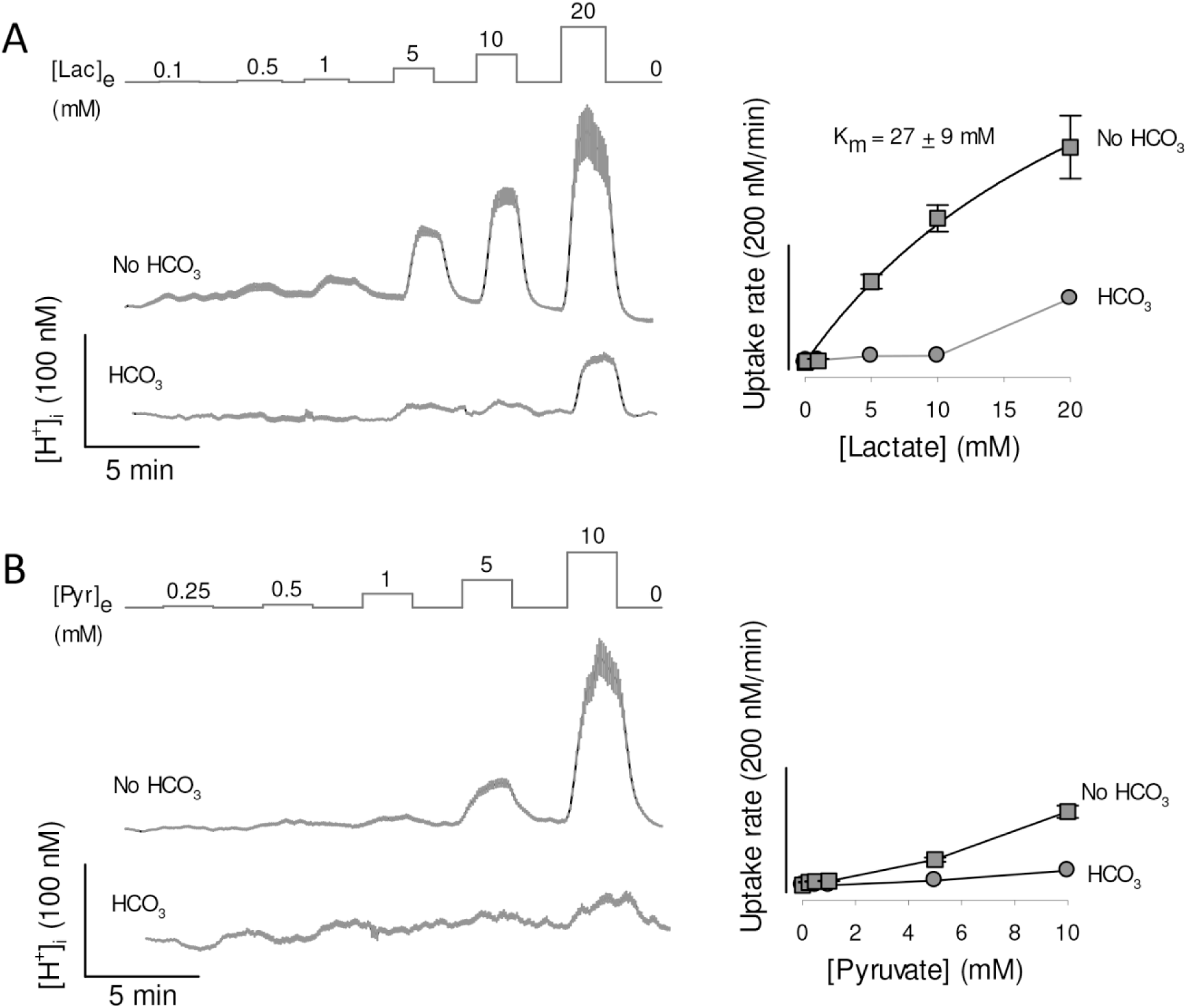
Lactate-and pyruvate-induced acidification in MDA-MB-231 cells. MDA-MB-231 cells were loaded with the pH-sensitive probe BCECF, which was calibrated as described in Methods. Resting proton concentration ranged between 36 to 45 nM (pH 7.35 to 7.44). (A) Cells were exposed to increasing concentrations of lactate, from 0.1 to 20 mM, in presence and absence of 24 mM HCO_3_ ^-^, equimolarly replaced by HEPES. Traces show intracellular proton concentration of 10 cells (mean ± SEM) in a single experiment, representative of three. Dose responses of the rates of acidification are shown in the right graph. The K_m_ was obtained by fitting a rectangular hyperbola to the data in the absence of bicarbonate. (B) Cells were exposed to increasing concentrations of pyruvate, from 0.25 to 10 mM, in presence and absence of 24 mM HCO_3_^-^, equimolarly replaced by HEPES. Traces show intracellular proton concentration of 10 cells (mean ± SEM) in a single experiment, representative of three. Dose responses of the rates of acidification are shown in the right graph.

### Recombinant MCT4 is also a high affinity lactate transporter

To explore the functional properties of recombinant MCT4, we used HEK293 cells. They possess abundant MCT1 ^18^ but it is still possible to use them to characterize a foreign transporter if the endogenous MCT1 is blocked pharmacologically, as recently demonstrated for the identification of a *Drosophila melanogaster* monocarboxylate carrier ^25^. As expected, the uptake of lactate by wild type HEK293 cells was blocked by AR-C155858 (Fig. 6A). Beyond our expectations, overexpression of MCT4 rendered the cells insensitive to the MCT1/2 blocker (Fig. 6B). We do not know how MCT4 overexpression suppressed the functionality of native MCT1 to such an extent, a phenomenon that may be of physiological interest, as MCT1 and MCT4 may co-exist in the same cells and use the same chaperone basigin/CD147 to reach the plasma membrane ^30^. A dominant role for MCT4 was confirmed by full inhibition of lactate uptake by diclofenac (Fig. 6B). Still, transport affinity was determined in the presence of AR-C155858 to ensure lack of MCT1 & MCT2 function. We found that HEK293-MCT4 cells transport lactate with a median K_m_ of 0.7 mM (Figs. 6C-E). Thus, high substrate affinity is also a property of recombinant MCT4.

**Figure 6.**
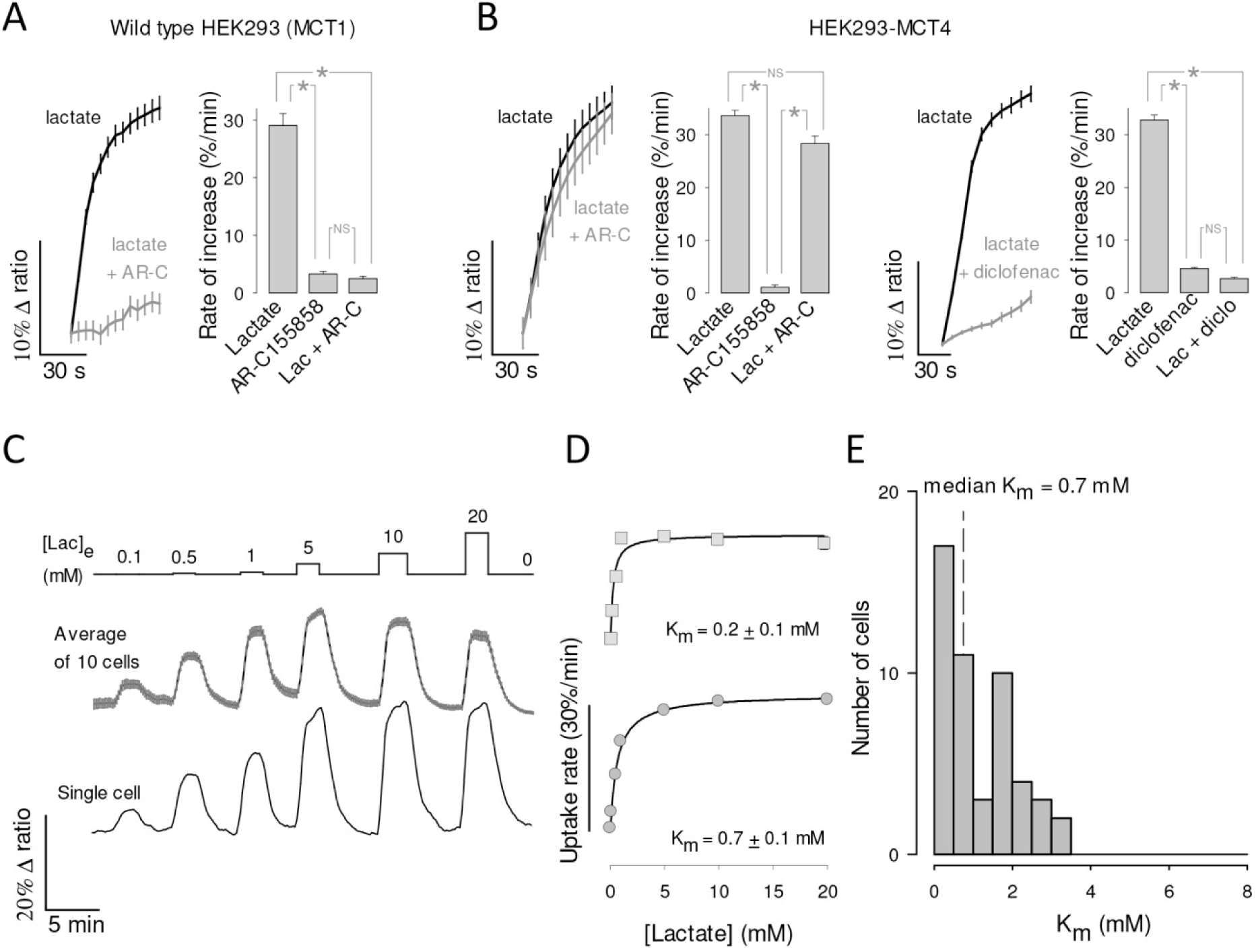
High affinity lactate transport in HEK293 cells overexpressing recombinant MCT4. (A) HEK293 cells expressing Laconic were exposed to 10 mM lactate in the presence and absence of 1 μM AR-C155858 (mean ± SEM of 10 cells). The bar graph summarizes the results of three experiments (mean ± SEM). (B) HEK293 cells co-expressing MCT4 and Laconic were exposed to 10 mM lactate in presence and absence of 1 μM AR-C155858 (left panel) or 1 mM diclofenac (right panel), mean ± SEM of 10 cells. Bar graphs summarize the results of three experiments (mean ± SEM). (C) HEK293 cells co-expressing MCT4 and Laconic were exposed to increasing concentrations of lactate, from 0.1 to 10 mM, as indicated. Responses of intracellular lactate in a single cell (bottom) and in 10 cells from a representative experiment (mean ± SEM, top) are shown. (D) Dose-response of the initial rate of lactate uptake, from the same cells depicted in C. K_m_ values were obtained by fitting a rectangular hyperbola to the data. (E) Frequency distribution from 50 cells in five experiments. Median K_m_ was 0.7 mM.

### High affinity MCT4-mediated lactate transport in human macrophages

The Warburg effect is important for the activation and operation of macrophages ^39, 40^, cells characterized by high MCT4 expression (Fig. 7A) ^41–43^. To study the affinity of MCT4 in these cells, monocytes were isolated from blood samples collected from healthy donors, transformed into macrophages *in vitro* and transduced with an adenoviral vector for Laconic. Experiments were carried out with undifferentiated macrophages (M0) and polarized macrophages (M1). In both developmental stages, the uptake of lactate was strongly inhibited by diclofenac but not by AR-C155858, evidencing a preferential role for MCT4 (Fig. 7B). K_m_ values were similar for M0 and M1 macrophages (Fig. 7C), with a pooled average of 1.2 mM.

**Figure 7.**
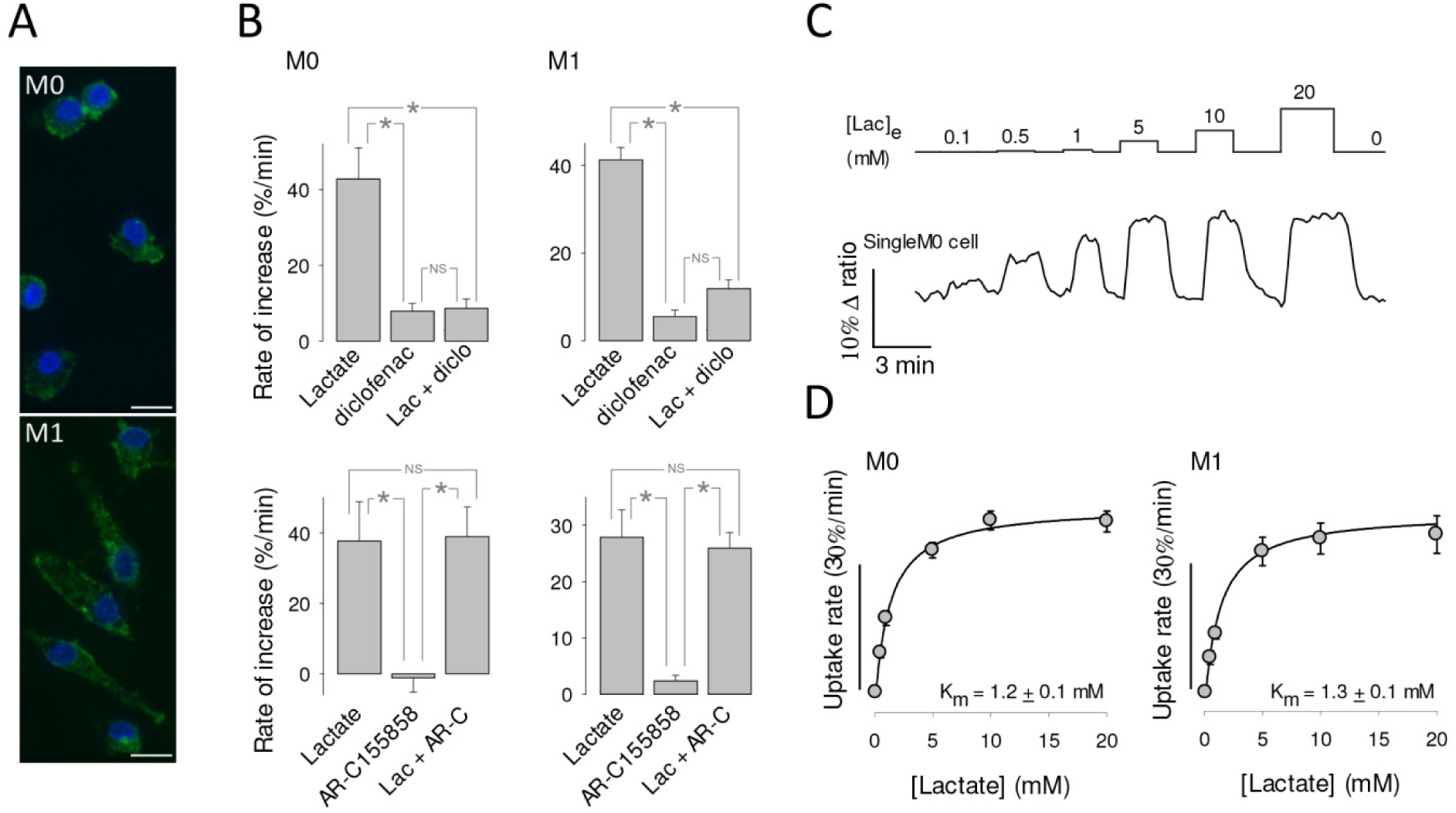
High affinity lactate transport in human macrophages. (A) Undifferentiated (M0) and polarized (M1) macrophages immunostained for MCT4 (green). DAPI-stained nuclei are shown in blue. Bar represents 10 μm. (B) The uptake of 1 mM lactate by macrophages was monitored before and during exposure to MCT inhibitors diclofenac (0.5 mM; diclo) or AR-C155858 (1 μM; AR-C). Bar graphs show the initial rates of uptake and the rate of accumulation elicited by the inhibitor itself. (C) Macrophages were exposed to increasing concentrations of lactate, from 0.1 to 20 mM, as indicated. The trace shows the response of an individual M0 macrophage. (D) K_m_ values were obtained by fitting a rectangular hyperbola to the data. Mean ± SEM of 11 cells in three experiments (M0) and 10 cells in three experiments (M1). A pool of M0 and M1 gave a K_m_ of 1.2 ± 0.1 mM (21 cells in six experiments).

### MCT4 but not MCT1 or MCT2 can export lactate against high ambient lactate

The impact of MCT isoforms on cellular lactate and pyruvate dynamics was gauged using numerical simulation based on the alternating conformer model of the transporter (Fig. 8A) ^34, 44^. The behaviors of MCT4 and MCT1 were first compared at physiological levels of lactate and pyruvate (Fig. 8B, left panel). Glycolytic cells were simulated by tuning mitochondrial pyruvate consumption and transporter dosage so that lactate was exported at 95% of the glycolytic flux ^5, 45^. For both isoforms there was pyruvate uptake. It seems remarkable that MCT4 imports almost as much pyruvate as MCT1, despite having an affinity eight times lower. This can be explained by a higher availability of the outward-facing carrier (T_IN_ in Fig. 8A), pushed by lactate on its way out. This pyruvate uptake helps to replenish the intracellular pyruvate pool and thus sustain lactate efflux, which otherwise would be capped at 90% of the glycolytic flux. The steady-state concentration of lactate and pyruvate were slightly higher in MCT1 cells, but on the whole both isoforms behaved similarly when simulated at low extracellular lactate (Fig. 8B, left panel).

**Figure 8.**
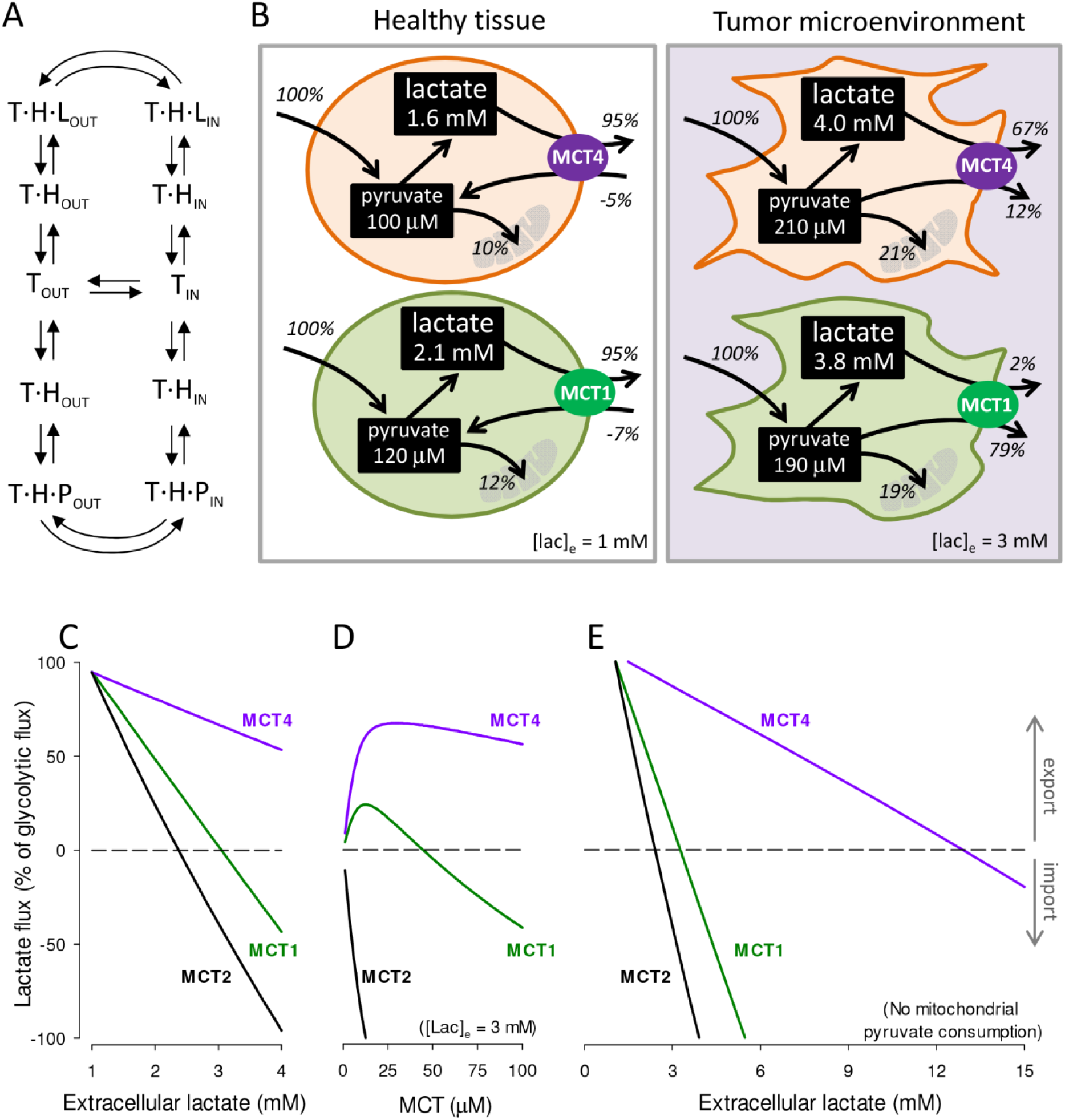
MCT4 is capable of lactate release in high lactate microenvironments. (A) Alternating conformer model of MCTs. The transporter (T) binds a proton (H) before binding either lactate (L) or pyruvate (P). Only empty and fully-loaded transporters alternate between outward-facing (out) and inward-facing (in) conformations. (B) Simulation of highly glycolytic cells. Glycolytic flux was fixed at 10 μM/s (100%). Rate constants were 0.01 s^-1^ (mitochondrial pyruvate import), 0.5 s^-1^ (LDH forward) and 0.025 s^-1^ (LDH reverse). Transporter quantities were 40 μM (MCT4), 42 μM (MCT1) and 3.8 μM (MCT2). Dynamics were simulated as specified in Methods. Extracellular lactate was 1 mM (left panel, healthy tissue) or 3 mM (right panel, tumor microenvironment). Extracellular pyruvate was 0.1 mM for both conditions. Fluxes are given as percentage of the glycolytic flux. (C) Effect of increasing extracellular lactate on lactate flux through MCT1, MCT2 and MCT4, starting from the conditions in B, left panel (healthy tissue). (D) Effect of increasing transporter dosage on lactate flux through MCT1, MCT2 and MCT4 at 3 mM extracellular lactate (tumor microenvironment). (E) Effect of increasing extracellular lactate on lactate flux through MCT1, MCT2 and MCT4 in the absence of mitochondrial pyruvate influx.

At elevated ambient lactate, such as is observed within tumors and inflammatory sites, a marked functional divergence between MCT4 and MCT1 became evident (Fig. 8B, right panel). Here MCT1-bearing cells became pyruvate producers while MCT4 cells maintained their lactate producing role and generated little pyruvate. The divergence was more marked at higher lactate levels (Fig. 8C) and at higher transporter dosages (Fig. 8D). With mitochondria unable to consume pyruvate, as would occur during in hypoxia, MCT1 cells reverted from lactate producers to consumers at 3.5 mM extracellular lactate whereas MCT4 cells reverted at 13 mM lactate (Fig. 8E). MCT2-bearing cells showed a strong tendency towards lactate consumption (Fig. 8D-F) consistent with the expression of this isoform in highly oxidative cells like neurons ^46^. For simplicity, the cellular NADH/NAD^+^ ratio in these simulations was fixed, that is, it was implicitly assumed that mitochondria compensate for deficits in NAD^+^ recycling at LDH. If this were the case, MCT4 cells will not only release more lactate than MCT1 cells will use less oxygen. Lactate and pyruvate fluxes are not only determined by MCTs, but also by glycolytic and mitochondrial fluxes and the redox ratio. Thus, these simulations do not cover every possible condition, but serve to demonstrate that all things being equal, MCT4 is more suited for lactate export than MCT1 and MCT2 at high ambient lactate levels.

## DISCUSSION

Our main conclusion is that MCT4 is a high affinity lactate transporter and that has a relevant affinity for pyruvate. A similar K_m_ for lactate of around 1 mM was determined in three different cell types including endogenous and recombinant MCT4, which suggests that this is a general property of the isoform. High affinity for lactate and a somewhat lower affinity for pyruvate confer MCT4-expressing cells the ability to export lactate against high ambient lactate levels, a role that is not possible for either MCT1 or MCT2, which cannot help losing pyruvate. This ability helps to explain why MCT4 is preferentially expressed in metastatic tumors, rapidly proliferating cells and hypoxic tissues.

How come MCT4 has been considered to be a low affinity lactate transporter with negligible affinity for pyruvate? When BCECF was first used to monitor monocarboxylate transport in 1990, bicarbonate was purposely omitted from experimental solutions “*to minimize intracellular buffering in order to produce greater and faster pHi changes when small amounts of lactate were introduced*” ^16^. Shortly afterwards, BCECF was used to estimate kinetic parameters, also in bicarbonate-free conditions ^17^. We confirm here that bicarbonate makes a big difference in the acidification induced by lactate. However, bicarbonate omission is not enough to eliminate the problem of buffering, because bicarbonate represents only a 30-50% of the buffering power of mammalian cells, the remainder being shared by protonable aminoacid residues, phospholipids, metabolites, etc ^36, 37^. As well as buffering, mammalian cells possess efficient systems for the extrusion of protons, including carbonic anhydrase, Na^+^/H^+^ exchangers and Na^+^/bicarbonate cotransporters, some of which are strategically located in the vicinity of MCTs ^38^. Of note, NBCe1 remains active even in the nominal absence of bicarbonate ^47^. A study in *Xenopus laevis* oocytes showed that the MCT4 activity is enhanced by membrane-anchored carbonic anhydrase. Significantly for affinity estimations, the effect of carbonic anhydrase was stronger at low lactate concentrations ^48^. Our interpretation of the bias introduced by pH measurements is that on the whole, the pH regulatory system is saturable. Challenged by low lactate loads, it copes well so that intracellular pH remains stable in spite of lactate influx. At higher lactate loads, the regulatory system is overwhelmed and cells acidify. In the presence of bicarbonate, pH regulation is even stronger, so that MCT-mediated pH changes are difficult to detect even at high lactate loads, and particularly in response to pyruvate, which is a less efficient substrate. The confounding effect of pH regulation leads to a biased estimation of affinity. Whereas proton buffering and muffling explain the high apparent K_m_ values previously reported for MCT4 in mammalian cells, it is not clear to us why a study based on radiolabeled lactate also reported a high K_m_ in MCT4-expressing oocytes (34 mM ^14^). The affinity measured here in HEK293-MCT4 cells suggest that over-expression may not account for the discrepancy, neither would genetic variability, because the splice variants of MCT4 do not include the protein coding region ^34^. Perhaps factors present in mammalian cells but not in *Xenopus* oocytes endow MCT4 with high affinity? Prime candidates are carbonic anhydrase and proton extrusion mechanisms, which when co-expressed in oocytes enhance the uptake of lactate ^38, 48^. Alternatively, the estimation of K_m_ in millimeter-sized oocytes may have been affected by unstirred layers that are not present in micrometer-sized mammalian cells, as discussed previously ^13^. A non-exclusive possibility is metabolism. The radiolabeled assay involved incubation for 20 minutes, during which some lactate may have been metabolized, an effect that would be more evident at low lactate loads. *Xenopus laevis* oocytes have a strong oxidative phosphorylation relative to glycolysis, producing CO_2_ from pyruvate 80-140 times faster than from glucose ^49^. They also have endogenous MCT and LDH ^50, 51^, and are therefore equipped to metabolize lactate. In the case of mammalian cells, oxidative phosphorylation is much slower than MCT-mediated transport, so it should not interfere significantly with the uptake assay. Still, a sizable metabolic interference in mammalian cells, a possibility that we do not favor, would mean that the affinity of MCT4 for its substrates is even higher than reported here. It has been proposed that MCT4 has a higher transport capacity than MCT1 ^14^. Calibrated lactate and pyruvate measurements accompanied by parallel measurement of MCT surface expression are needed to address the pending question of transport capacity.

### MCT4 versus MCT1 and MCT2

The most widely expressed, house-keeping member of the monocarboxylate transporter family is MCT1. It has an affinity for pyruvate 5-10 times higher than that for lactate, commensurate with the ratio between the physiological concentrations of the two substrates. MCT1 plays a major role in whole-body energy homeostasis and in the distribution of redox potential between organs ^52^. It is also widely expressed in tumors, where it mediates both the export and import of lactate ^12, 53^. Hypoxic cells may only use glycolysis to generate ATP, and to sustain glycolysis they need to recycle the NADH produced at GAPDH. With oxidative phosphorylation disabled in the absence of oxygen, NADH may only be recycled at LDH, the enzyme that converts pyruvate into lactate. Thus, to generate energy, hypoxic cells need to release lactate but not pyruvate, a task that is fitting for MCT4, but not for MCT1. This helps to explain why only the expression of the MCT4 is under the control of HIF-1α ^11^. At variance with hypoxic cells, which require glycolytic ATP production for survival, cancer cells are capable of generating their ATP in mitochondria by oxidative phosphorylation ^45^. For reasons that are not fully understood, but which include the interstitial effects of lactate and protons ^5^, some cancer cells engage in a glycolytic frenzy to export almost every glycolytic carbon in the form of lactate ^1–4^. They do this against elevated ambient lactate levels caused by inflammation, hypoxia and/or the Warburg effect in neighboring cancer and stroma cells ^4–6, 9^. According to our numerical simulations MCT1 cells may only produce lactate at low ambient lactate levels, because at high lactate levels they cannot avoid producing pyruvate. Consistently, pharmacological MCT1 inhibition in breast cancer cells was found to inhibit the release of pyruvate but not that of lactate ^54^. In contrast, MCT4 exports lactate regardless of extracellular lactate levels.

As well as to contributing to the understanding of high-lactate microenvironments, the revised kinetic properties we report here may inform the development of urgently-needed specific MCT4 blockers.

## ACKNOWLEDGMENTS

We thank Karen Everett for critical reading of the manuscript and José Sarmiento (Universidad Austral de Chile) for help with confocal microscopy of MCT4 in human macrophages. This work was partly supported by Fondecyt grants 11150930 to ASM and 1160317 to LFB. The Centro de Estudios Científicos (CECs) is funded by the Chilean Government through the Centers of Excellence Basal Financing Program of CONICYT.

